# *Capsicum chinense* Jacq. derived glutaredoxin (*CcGRXS12*) alters phytohormonal pathways and redox status of the cells to confer resistance against pepper mild mottle virus (PMMoV-I)

**DOI:** 10.1101/2023.02.01.526735

**Authors:** R. M. Saravana Kumar, S.V. Ramesh, Z. Sun, Sugitha Thankappan, Asish Kanakaraj Binodh

## Abstract

Glutaredoxins (Grxs) are small, ubiquitous, multi-functional proteins present in different compartments of plant cells. A chloroplast targeted class I GRX (*CcGRXS12*) gene was isolated from *Capsicum chinense* during the pepper mild mottle virus (PMMoV) infection. Functional characterization of the gene was performed in *N. benthamiana* transgenic plants transformed with native *C. chinense GRX* (*Nb:GRX*), *GRX*-fused with GFP (*Nb:GRX-GFP*) and *GRX* truncated for the chloroplast targeting sequences but fused with GFP (*Nb*:Δ*2MGRX-GFP*). Over-expression of *CcGRXS12* inhibits the PMMoV-I accumulation at late stage of infection and is accompanied with the activation of SA- pathway pathogenesis related (PR) transcripts, and suppression of JA/ET- pathway transcripts. Further the reduced accumulation of auxin-induced Glutathione-S-Transferase (pCNT103) in *CcGRXS12* over expressing lines indicates that the protein could able to protect the plants from the oxidative stress caused by the virus. PMMoV-I infection increases accumulation of pyridine nucleotides (PNs) mainly due to the reduced form of PNs (NAD(P)H) and it was higher in *Nb:GRX-GFP* lines compared to other lines where infection is limited. Apart from biotic stress, *CcGRXS12* protects the plants from abiotic stress conditions caused by H_2_O_2_ and herbicide paraquat. CcGRXS12 exhibits GSH-disulphide oxidoreductase activity *in vitro* however devoid of complementary Fe-S cluster assembly mechanism in yeast.

## 1. Introduction

Plant virus invasion subjugates the host machinery to express viral genes and induces oxidative stress conditions (Wang et al., 2021; Akbar et al., 2020). Plants manifest different strategies to arrest the spread of virus *viz*: development of hypersensitive response (HR) (Balint-Kurti, 2019); *R*- gene mediated resistance (Palukaitis & Yoon 2020); silencing of viral genes (Wang et al., 2012; Ismayil et al., 2018) and RNA decay (Li and Wang, 2019). Apart from *R*-gene mediated resistance, many other plant-derived genes also protect the plants from the oxidative damage caused by virus through scavenging the ROS accumulation. ROS is scavenged through enzymatic proteins such as superoxide dismutase (SOD), catalase (CAT), peroxidase (PRX), ascorbate peroxidase (APX), glutathione S-transferase (GST), glutaredoxin (GRX) and glutathione peroxidase (GPX) (Waszczak et al., 2018). Exploring the functions of the differentially expressed host genes during the plant-pathogen interaction provides a better comprehension of plant genes-mediated resistance and also helps in designing a novel plant-protection strategy (Marmonier et al., 2022). In this study, Pepper Mild Mottle virus-Italian strain (PMMoV-I) belonging to *Tobamovirus* genus causing a serious economic losses to pepper crops (*Capsicum chinense*) (Wetter et al., 1984) was utilized to analyze the role of host-derived genes against virus infection. Through mRNA differential display PCR and RACE-PCR, a cDNA fragment corresponding to class I Glutaredoxin (*CcGRXS12*) gene was isolated from *C.chinense* plants during the compatible (PMMoV-I) and incompatible (PMMoV-S) viral infection.

Glutaredoxins (GRXs) are small, ubiquitous, low molecular weight oxido-reductases sharing the structure with thioredoxin family of proteins and are well conserved among prokaryotes and eukaryotes (Holmgren, 1995). The number of GRXs reported in the photosynthetic organisms is found to be high and are classified into six classes. Class I (C1, C2, C3, C4, C5/S12 subgroup), Class II (S14, S15, S16, S17) and Class III (21 CC-type in Arabidopsis) are relatively well-characterized while other GRXs require further analysis (Couturier et al., 2009a). Individual class of GRX proteins has different catalytic activities and specific functions and thus various plant-derived GRXs coordinate different functions. Class I GRXs reduce the protein-protein disulphide bonds and protein-glutathione disulfide bonds by utilizing GSH as their reducing equivalent supplier (Rouhier et al., 2007). Class II GRXs reduce protein-glutathione disulphide bonds by utilizing ferredoxin-thioredoxin reductase (FTR) as their reducing equivalent supplier (Zaffagnini et al., 2008). Biochemical characterization of the Class III CC-type GRX proteins remain elusive as the soluble form of the protein could not be produced in the bacterial system (Couturier et al., 2010). The oxidoreductase property of the GRXs enables the protein to take part in many redox-dependent pathways leads to various protein activation/deactivativation.

Recent studies have divulged the importance of GRXs during plant-pathogen interactions. GRXs cause susceptibility of the plants to the necrotrophic pathogens while the effect over biotrophic pathogens is quite different. Salicylic acid (SA) induced CC-type GRXs (GRXC9/ROXY19 and GRXS13/ROXY18) cause susceptibility to the necrotrophic pathogen *B. cinerea* (Ndamukong et al., 2007; La camera et al., 2011) by over accumulating H_2_O_2_. Similarly, over expression of rice and Arabidopsis CC-type glutaredoxin (ROXY1, ROXY2) accumulate H_2_O_2_ and enhances the plants’ susceptibility to *B. cinerea* (Wang et al., 2009). Yang et al., (2022) have shown that over expression of *GsGRX4* makes the plants susceptible to *B.cinerea* by accumulating higher H_2_O_2_ and also by suppressing the JA content and JA related marker gene. A member of the tomato CC-type GRX (*SlGRXC6*) reduce the accumulation of tomato leaf curl virus (TLCV) by interacting with the virus protein (Zhao et al., 2021). Over expression of the rice Class I GRX (*OsGRX20*) mediates resistance against bacterial blight (*Xanthomonas oryzae* pv. Oryzae; Xoo) (Ning et al., 2018). Tomato Class II glutaredoxin (*SlGRX1*) has no role against plant virus (Guo et al., 2010).

Phytohormonal pathway activation plays major roles during the pathogen attack and also at the time of plant growth & developments (Ma & Ma., 2016; Bozbuga et al., 2022). During pathogen attack, systemic acquired resistance (SAR) is developed at the distal part of the infection site, concomitant with the activation of various phytohormome pathways and accumulation of the corresponding pathogenesis related (PR) proteins (Van Loon and Van Strien, 1999). In general, resistance against biotrophs is mediated through SA-pathway activation, while jasmonic acid/ethylene (JA/ET)-pathway acts against necrotrophs. Plants have the ability to regulate the phytohormonal pathway based on the type of pathogen it encounters. SA pathway activation during the pathogen attack, suppresses the JA and auxin-dependent defence pathways (Vlot et al., 2009; Yang et al., 2015; Yuan et al., 2017) to fine tune the defense reaction. In plants, different phytohormones induce the expression of different GRXs (Yang et al., 2021; Sharma et al., 2013; Herrera-Vásquez et al., 2015; El-Kereamy et al., 2015; Malik et al., 2020) suggesting that GRXs transmit the information of phytohormonal pathway activation.

In the SA/JA signal transduction pathway, many transcription factors (TGA, ORA59) and various proteins are involved and GRX activates/deactivates the signaling components through its oxido-reductase property. In the NPR1-TGA transducing system, reduction of the co-activator (NPR1) and the transcription factor (TGA) is occurred before to their interaction. Following the NPR1-TGA interaction, the TGA protein could bind the as-1 elements of PR gene’s promoters and mediates the transcription of PR genes (Després et al., 2000). SA- inducible GRX480 suppress the expression of JA induced PDF1.2 by interacting with TGA2 factor (Ndamukong et al. 2007). In Arabidopsis, 17 of the 21 CC-type GRXs interact with TGA2 factor (Zander et al., 2012) implying GRXs interaction with TGA is an inevitable process. Apart from pathogen attack, many developmental activities are also influenced by the interaction of TGA transcription factors and GRXs (Gutsche et al., 2017; Ehrary et al., 2020; Ruan et al., 2018, 2022; Li et al., 2019; Uhrig et al., 2017). GRXs activate the TGA factor through their post-translational modification (Hou et al., 2019). SA pathway activation inhibits the expression of JA/ET-induced genes, through the repression of ORA59 (Pre et al., 2008; Van der Does et al., 2013) and CC-type GRXs are reported to suppress the ORA59 activation (Zander et al., 2012). Apart from acting as transducing element, GRXs influence the phytohormones pathways through synthesizing them (El-Kereamy et al., 2015). GRXs are actively engaged in the Fe-S cluster assembly mechanism (rev Lu, 2018).

In the present study, the functional characterization of *CcGRXS12* gene was carried out by over-expressing in *N.benthamiana* domin model plants. These plants are susceptible to PMMoV-I viral infection and the plants get recovered from PMMoV-I infection at later stage. This recovery phenomenon helps in obtaining molecular insights regarding the resistant mechanisms conferred by the gene during plant-virus interaction. The indispensable roles of GRX in the activation of phytohormone pathways were studied by analysing the corresponding PR transcript(s) accumulation. *CcGRXS12* role in altering the redox status of the cell was studied by analysing the accumulation of oxidized and reduced forms of pyridine nucleotides (PNs). In addition, we have deduced the function of *CcGRXS12* in relation to plant’s abiotic stress tolerance and also in Fe-S cluster assembly mechanism.

## 2. Materials and Methods

### 2.1. Plant materials and virus inoculation

*C. chinense* N.J. Jacq. PI159236 (*L^3^L^3^*) and various *N. benthamiana* transgenic plants were maintained in growth chambers at 32°C with 16 hrs of photoperiod, light intensity of 8000 lx and 70% relative humidity. For viral inoculation, first pair of the developed leaves from the plants was mechanically inoculated with purified virions at a concentration of 50μg/mL in inoculation buffer (0.02 M sodium phosphate buffer, pH 7.0), using carborundum as abrasive material. At 7 days post inoculation (dpi), samples were taken from inoculated and systemic leaves, for 14 and 21 dpi, samples from systemic leaves and for 28 dpi, samples were collected from the asymptomatic (recovered leaves) along with the symptomatic leaves.

### 2.2. Viral strain, purification, and viral RNA extraction

Pepper mild mottle virus – Italian strain (PMMoV-I), reporter earlier, was used in this work (Wetter et al., 1984). The protocols for purification of virion and viral RNA extraction were as enumerated earlier (Alonso et al., 1991; García-Luque et al., 1990).

### 2.3. Isolation of *CcGRXS12* and sequence analysis

Previous work in our lab, characterised the *Capsicum chinense* (*L^3^L^3^*) (PI 159236)-derived transcript corresponding to Glutaredoxin gene when the plants were infected with compatible (PMMoV-I) and incompatible (PMMoV-S) viral strains. The complete sequence of the gene was characterised by following “mRNA Differential Display PCR” (Liang and Pardee, 1992) and RACE-PCR (Chenchik et al., 1998.) techniques. The chloroplast targeting region of the protein was predicted by chloroP 1.1 programme (Emanuelsson et al., 1999). Protein sequence alignment was performed by Clustal W program (http://www.ebi.ac.uk/Tools/msa/clustalw2/).

### 2.4. Measurement of oxidoreductase activity

The purified native protein was used to determine the GSH-disulfide oxidoreductase activity by HED assay (Holmgren and Aslund, 1995). To express the protein in prokaryotic system, the truncated CcGRXS12 protein (63 amino acids in length) tagged with 6X His-tag at its N-terminal region was cloned in pQE-1 vector (obtained from Dra.Maria Teresa). *E.coli* M15 (pREP4) strain was transformed with the plasmid harboring the cloned gene construct. Protein expression and native purification was performed by using His-select Nickel affinity gel resin column (Sigma Aldrich, USA) according to the manufacture’s instruction. GRX activity was measured as an oxidation of NADPH in a reaction comprising 1mM GSH, 0.7mM β-hydroxy ethyl disulphide (HED), 0.25 mM NADPH and 6.4μg/mL glutathione reductase in Tris-Cl pH 7.4. The reaction mixture was incubated at room temperature for 2 min then the decrease in OD at 340 nm was recorded in a spectromax micro-plate reader for 1 min at room temperature. His-GRX protein was added to the same cuvette and the decrease in absorbance at 340 nm was recorded. Measured activities were normalized by correcting for the absorbance before the addition of GRX protein. One unit of activity is defined as the consumption of 1μmol of NADPH per minute calculated from the expression (ΔA340×V)/ (min× 6.2), where V is the cuvette volume in mL and 6.2 is the mM extinction coefficient for NADPH. Three independent experiments were performed at each substrate concentration, and the apparent *K_m_* value and *K_cat_* values were calculated by non-linear regression using the program SigmaPlot 12.0.

### 2.5. Transgenic plants

*Nicotiana benthamiana* Domin transgenic plants constitutively expressing GFP (*Nb:GFP*; line 11), full length GRX (*Nb:GRX*; line 3), full length GRX fused with GFP (*Nb:GRX-GFP*; line 16) and truncated GRX (63aa) fused with GFP (*Nb*:Δ*2MGRX-GFP*; line 40) were used. These transgenic lines were provided by Dr. Maria Teresa Serra Yoldi (Montes-Casado et al., 2010). The constructs were driven at their N-terminal by CaMV35S promoter and have NOS terminator at its C-termini.

### 2.6. Plant total protein extraction

Total protein from the fresh leaf samples (1mg) were extracted in 5 μL Laemmli buffer (Laemmli, 1970). Briefly, samples were heated to 95°C for 5 min followed by sonication for 5 min in a water sonicator and then clarified by centrifugation at 20,000 g for 5 min in a microcentrifuge. Total protein extracts (5 μL) were resolved in SDS polyacrylamide gels (SDS-PAGE) using 17.5% and 4.5% polyacrylamide as solving and concentrating gels, respectively according to Laemmli (1970).

### 2.7. Viral coat protein analysis

Total plant protein extracts (5 μl) and a viral coat protein extracts of 5, 2.5 and 1.25 ng were electrophoresed on 17.5% and 4.5% SDS-PAGE gel. The proteins were stained with coomassie blue R250 and PMMoV-I coat protein was quantified using a densitometer and the quantity one software (Bio-Rad, Hercules, CA, USA).

### 2.8. Immunoblot analysis

Total proteins separated by SDS-PAGE were electrotransferred onto PVDF membranes (Amersham). The membranes were initially blocked with PBST (3.2 mM Na_2_HPO_4_, 0.5 mM KH_2_PO_4_, 1.3 mM KCl, 135 mM NaCl, 0.05% Tween 20, pH 7.2) containing 5% skimmed milk for 30 min. For the immunodetection of GRX, GFP and viral Coat protein (CP), the electro transferred membranes were incubated overnight at 4°C with the specific antisera of His-GRX (1/1000; raised in our lab), GFP specific polyclonal antibody (1/250; Santa Cruz Biotechnology, INC.) and PMMoV-I CP (1/1000; raised in our lab). Detection of antigen-antibody complexes was carried out with peroxidase-conjugated goat anti-rabbit IgG (GARPO) (Nordic) at 1/5000 dilution. The immunoreaction was visualized with ECL chemiluminescence kit (GE Healthcare Amersham) following manufacturer’s instructions. The enzymatic reaction produces a luminescent compound that is detected by visible light sensitive films (Hyperfilm, Pharmacia).

### 2.9. Protoplast infection assay

Protoplasts were obtained from the four different *N.benthamiana* transgenic lines as described by Ruiz del Pino et al., (2003) with a minor change in protocol. Protoplasts were washed a couple of times with ice-cold solution of 12% mannitol containing 6mM CaCl_2_. Protoplasts were counted and adjusted to the concentration of 1.3×10^6^ protoplasts/mL using ice-cold electroporation buffer (12% mannitol, 6 mM CaCl_2_, 80 mM KCl, pH 7.2). For protoplasts infection, 4×10^5^ protoplasts in a final volume of 300μL were taken in a 0.2 cm electroporation cuvette along with 20 μg of PMMoV-I RNA and kept on ice. For transformation, a single pulse of 0.12kV and 125 μF was applied with a Gene Pulser (Bio-Rad laboratories, Hercules, CA) immediately after addition of 20 μg of viral RNA. Following the pulse, the protoplasts were kept on ice for 20 min and diluted in CPW-M13. Later, centrifuged at 80×g for 5 min and diluted to 5×10^5^ protoplasts/mL in CPW-M13 and incubated at 25°C under dark. Protoplasts were harvested at 16, 24 and 48 hrs with a short spin of 134g, and a couple of washing in mannitol buffer and the resultant protoplast was re-suspended in 5X Lammeli buffer. The viral infection was detected by western blot method using PMMoV-I coat protein specific antibody.

### 2.10. Sub-cellular localization studies of CcGRXS12 protein

For analyzing the sub cellular localization of CcGRXS12 protein, protoplasts obtained from the different *GFP* expressing transgenic lines (*Nb:GFP*, *Nb:GRX-GFP* and *Nb*:Δ*2MGRX-GFP*) were fixed with 4% formaldehyde in 9% mannitol (pH 7.2). To label the nuclei, DAPI staining was performed. In brief, 10^4^ protoplasts were incubated with 100μL of 2μg/mL DAPI solution in PBS-9% mannitol for 5 min. To that 900μl of PBS-9% mannitol was added and kept for another 15 minutes, and then the protoplasts were collected and washed with PBS-9% mannitol for twice and finally suspended in 100 μL PBS-9% mannitol. Around 20μL of this sample was loaded over poly-L-lysine coated glass slides for visualization. For compartmentalization study, autofluorescence from chlorophyll and DAPI staining at nuclei were analyzed. Different fluorescent signals were detected at specific wavelengths [GFP detection at 550nm (excitation 580nm), chlorophyll autofluoresence at 540nm (excitation 600nm) and DAPI at 610nm (excitation at 650nm)] using a TCSP Leica microscope.

### 2.11. RNA isolation and probe preparation

Total leaf RNA was extracted according to the method prescribed by Chomcynski and Sacchi, (1987) by using TRIzol reagent and following the manufacturer’s instruction with a slight modification Briefly, additional centrifugation at 12,000 g for 10 min at 4°C after plant tissue homogenization and an additional final RNA precipitation step with 0.3 M sodium acetate pH 5.2 and 2.5 volumes of 100% ice-cooled ethanol were performed for overnight. The digoxygenin-labelled RNA probes were prepared using the linearized plasmids harbouring various gene products according to the instructions enumerated in MAXIscript kit manual. The cloned gene products, for preparing the riboprobes, were: salicylic acid pathway marker proteins (PR-1, PR-2a and PR-5), the JA/ET pathway marker protein (PR-2d) and the gene for auxin inducible Glutathione-S-Transferase (GST) (pCNT103) (obtained from Carmen Castresana), the clone pT-CPS containing the 593 bp from PMMoV-S CP (Gilardi et al., 1998)

### 2.12. Northern blot hybridization

For Northern blot analysis, RNA samples (10 μg) were denatured at 65°C for 4 min in MOPS-Acetate-EDTA buffer (20 mM MOPS, 15 mM sodium acetate, 3 mM EDTA pH 7.0) in the presence of 10% formamide, 0.9% formaldehyde and 0.06 mg/mL ethidium bromide. Then the samples were electrophoresed onto 1.5% agarose-formaldehyde gels containing MOPS buffer (50 mM MOPS, 0.4 mM EDTA pH 7) and 6% formaldehyde, using MOPS electrophoresis buffer under a current of 5 V/cm.

Once visualized by UV light illumination, the RNAs were transferred by capillarity to nylon membranes (Hybond-N, Pharmacia), as described in Sambrook et al., (1989), and fixed to the membrane by irradiation with UV light (120 mJ) in a UV Stratalinker apparatus (Cultek). The membranes with transferred RNA were stored in cold condition for later use.

For hybridization with digoxigenin-labelled RNA probes, the membranes were incubated with standard buffer (5xSSC, 0.1% sodium-lauroylsarcosine, 0.02% SDS and 2% blocking reagent, blocking reagent is provided by manufacturer) for 2 hrs at 65°C. Then the membranes were hybridized overnight at 68°C with specific RNA probes in standard buffer containing 100 ng of the corresponding probe. After hybridization, the membranes were washed twice in (2x SSC solution, 0.1% SDS) for 15 min each at room temperature (RT) and twice in (0.1 x SSC, 0.1% SDS) for 15 min at 65°C. Probe detection was performed using the DIG luminescent detection kit (Roche, Penzberg, Germany) according to manufacturer’s protocol. In brief, membranes were incubated in blocking solution of maleic buffer (0.1M Maleic acid, 0.15M NaCl) containing 1% blocking reagent- for 30 min, then in antidigoxigenin alkaline phosphatase conjugated antibody at 1:10,000 for 30 min followed by washing twice in washing buffer. After equilibration in detection buffer, membranes were incubated with chemiluminescent substrate CSPD and exposed to X-ray sensitive films (Hyperfilm, Pharmacia) for 30 min.

### 2.13. Measurement of pyridine nucleotide (PN) contents

PN contents in the mock and PMMoV-I-infected plants at 28 dpi were analyzed by grinding 30 mg of fresh leaf sample with ball bearings and centrifuged for 12,000 g for 1min. The supernatant obtained was used for selective extraction of NAD(P)H in acid medium and NAD(P)+ in alkaline medium (Hajirezaei et al., 2002). The samples were neutralized to the final pH of 8.0 to 8.5 and made as 100μL aliquots and frozen at −80°C for later analysis. PNs in the neutralized extracts were determined following Gibon et al., (2004). Statistical difference was analysed by SAS 9.1 programme (SAS 9.1 Inc.) using one-way analysis of variance (ANOVA).

### 2.14. Abiotic stress tolerance assay

Role of *CcGRXS12* in abiotic stress tolerance was analyzed by growing different *N. benthamiana* transgenic plants in one-half MS media in the presence and absence of 3mM H_2_O_2_ and 1 μM paraquat. The seedlings were grown horizontally for the first 3 days and then grown in vertical position for another one week. Root length was measured after 10 days. The experiments were performed in triplicates of 30 seedlings each and repeated thrice. The seedlings were grown at 25°C and 16 h of photoperiod. Statistical difference was analysed by SAS 9.1 programme (SAS 9.1 Inc.) using one-way analysis of variance (ANOVA).

### 2.15. Heterologous expression of *CcGRXS12* in *S.cerevisae*

To clone and express CcGRXS12 mature protein in yeast cells, pGEM-T easy vector (pGEM-GRX), harbouring the full length *CcGRXS12* gene was used. Gene specific primers were designed to amplify the full length gene excluding the chloroplast targeting region. Further in order to prevent codon breakage while expression in yeast, we introduced ‘g’ nucleotide (small letter) immediately after the *Not* I site in the forward primer (5’-ATAAGAATGCGGCCGCgTCGGGTTCATTCGGGTCC-3’) which introduce alanine at the end of the mitochondrial targeting region. The reverse primer (5’-GAAGATCTGCTTTCTGTTTTTCCAGGATTA-3’) had an overhang of *Bgl* II site. The amplified fragment was cloned in to pMM221 vector (obtained from Dr.E.Herrero, Universitat de Lleida, Lleida, Spain) which contains the yeast mitochondrial targeting sequence at its 5’ end and 3HA/His6 tag at its 3’ end (Molina et al., 2004). The resultant plasmid containing the *CcGRXS12* gene sequence was transformed into yeast Δ*grx5* mutants and the selection of the transformants were performed following Rodriguez-Manzaneque et al., (1999).

### 2.16. Growth conditions of *S.cerevisae*

The yeast strains employed in this study are: wild type (W303-1B; WT), Δ*grx5* mutant (MML100; Δ*grx5*) and yeast *GRX5* expressing in Δ*grx5* mutants (MML240; Δ*grx5/GRX5*) were already reported in Rodriguez-Manzaneque et al., (1999) and the *CcGRXS12* transformed in yeast Δ*grx5* mutant strain (Δ*grx5/CcGRXS12*) (in this work). All yeast strains were grown in YPD, respiratory defective YPG and SC media as mentioned in Rodriguez-Manzaneque et al., (2002). For oxidant sensitive assay, above mentioned yeast strains were grown in the media containing 0.3mM tert-butyl hydroperoxide (t-BOOH) and 1.25mM diamide (Sigma Aldrich, USA) in a serial dilution of 1:10 and incubated for 3 days.

### 2.17. *CcGRXS12* in Fe-S cluster assembly mechanism

Role of *CcGRXS12* in Fe-S cluster assembly mechanism was studied by analysing the accumulation level of free iron (Fish, 1988), and also by measuring the relative ratio of aconitase to malate dehydrogenase activity (Robinson et al., 1987) for the aforementioned yeast strains.

## 3. Results

### 3.1. Sequence features of *Capsicum chinense* glutaredoxin

Sequence analysis of *Capsicum chinense* glutaredoxin gene (*CcGRX)* showed that the ORF comprises 531bp and its amino acid prediction showed it codes for 177aa. Protein sequence analysis revealed that it possess CSYS active site which is the characteristics of Class I / S12 GRXs. Based on the analysis, the isolated gene was named as *CcGRXS12*. Sequence alignment of the isolated GRX with the well characterized Poplar GRX (PtGRXS12) (Coutruier et al., 2011) protein showed that apart from CSYS active site, other motifs such as ELD, TVPN, GG and second cysteine were conserved between these two proteins (Fig.1A).

**Fig.1.**
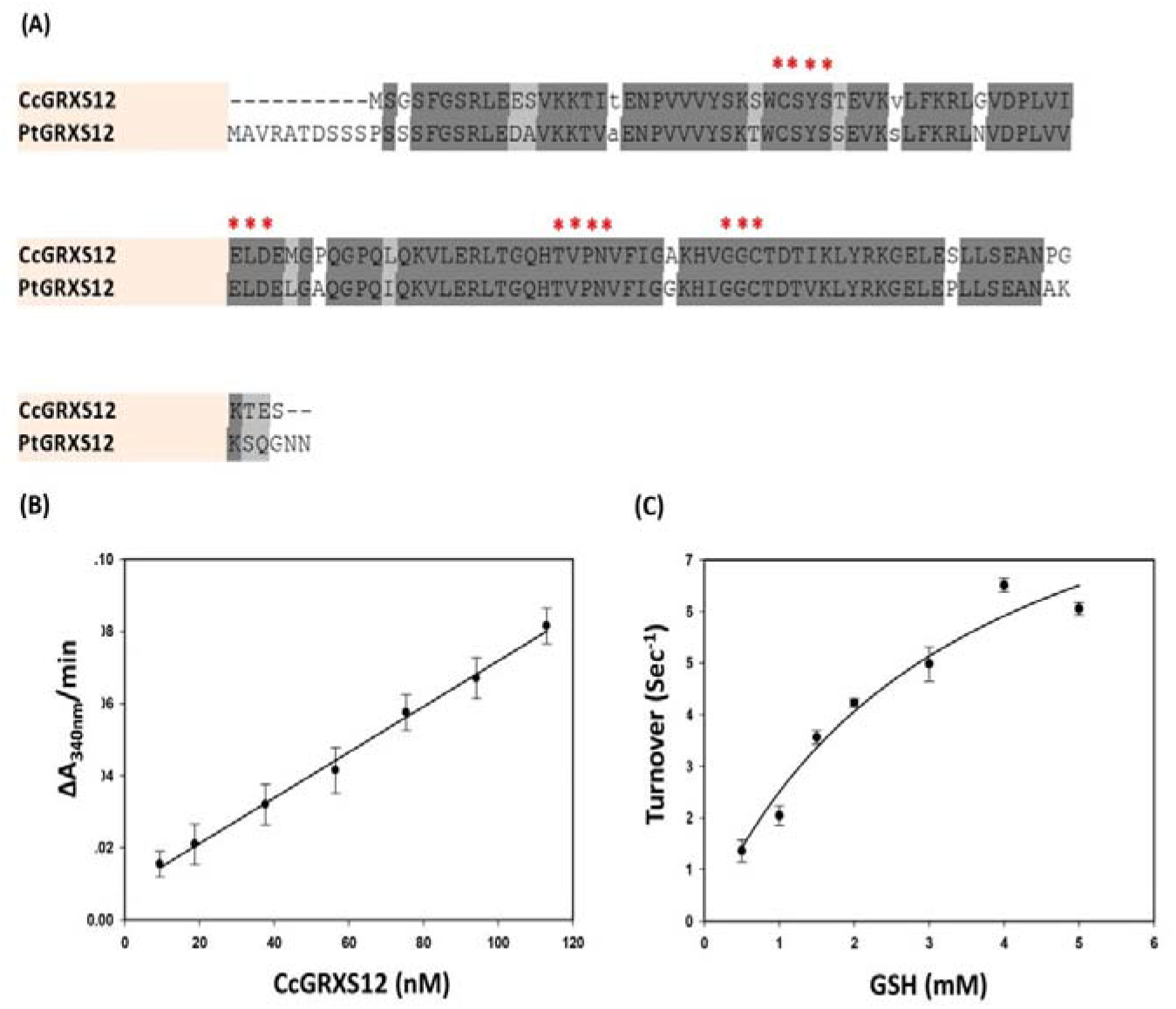
(A) Comparative protein sequence alignment of isolated GRX (CcGRXS12) with GRX characterized from Poplar (PtGRXS12). Sequence features have shown that besides the conserved CSYS active site, there are many conserved regions between these two proteins. (B) Linear dependence of HED activity on *Cc*GRXS12 concentration is expressed as Δ*A*_340_/min. (C). Variations of the apparent turnover during hydroxyethyl disulphide (HED) assay were calculated by varying concentrations of GSH (0.5 to 5mM). The data are represented as mean ± S.D. The best fit was obtained using the Michaelis-Menten’s equation using non-linear regression analysis.

### 3.2. CcGRXS12 possesses GSH-disulfide oxidoreductase activity

Mature form of the CcGRXS12 protein (114 aa) without the chloroplast targeting region was cloned in pQE1 vector and expressed in M15 (pREP4) *E.coli* expression system. Purification of native protein was carried out using nickel affinity column. The predicted molecular mass and pI for the protein were 13.8 kDa and 8.24 respectively. Specific activity of the protein (0.12U/nM) was calculated by varying the concentration of His-CcGRXS12 from 10 nM to 125 nM by keeping GSH concentration as 1 mM (Fig.1B). The correlation kinetics between the protein and GSH was measured by keeping the protein concentration at 84 nM and varying the GSH concentration from 0.5 mM to 5 mM (Fig.1C). The apparent *Km* (4.9 ± 1.9 mM) and apparent turnover *Kcat* (14.63 ± 4.8 sec^−1^) values were calculated using the Michaelis-Menten equation and the catalytic efficiency value (*K_cat_/K_m_* (*M^−1^Sec^−1^*)) was found to be 3.05×10^3^. *In vitro* assay showed that CcGRXS12 protein participates in deglutathionylation reaction.

### 3.3. Selection and analysis of CcGRXS12 over-expressing lines

For investigating the functional role of GRX, different *N.benthamaina* transgenic plants were used. The gene constructs developed and utilized for the genetic transformation of *N. benthamaina* are depicted (Fig.2A). The transgenic plants were grown in a media containing high concentration of kanamycin. The expression level of the gene in the T3 transgenic plants were analyzed by Western blot method (Fig. 2B). Analysis showed that the expression of *CcGRXS12* in *Nb:GRX-GFP* lines was found to be ~ 10 times higher than the other two *CcGRXS12*-expressing lines. Phenotypic analysis reveals that *Nb*:Δ*2MGRX-GFP* lines show stunted growth and wide leaf surface area when compared with free *GFP* and other *CcGRXS12* over-expressing lines.

**Fig.2.**
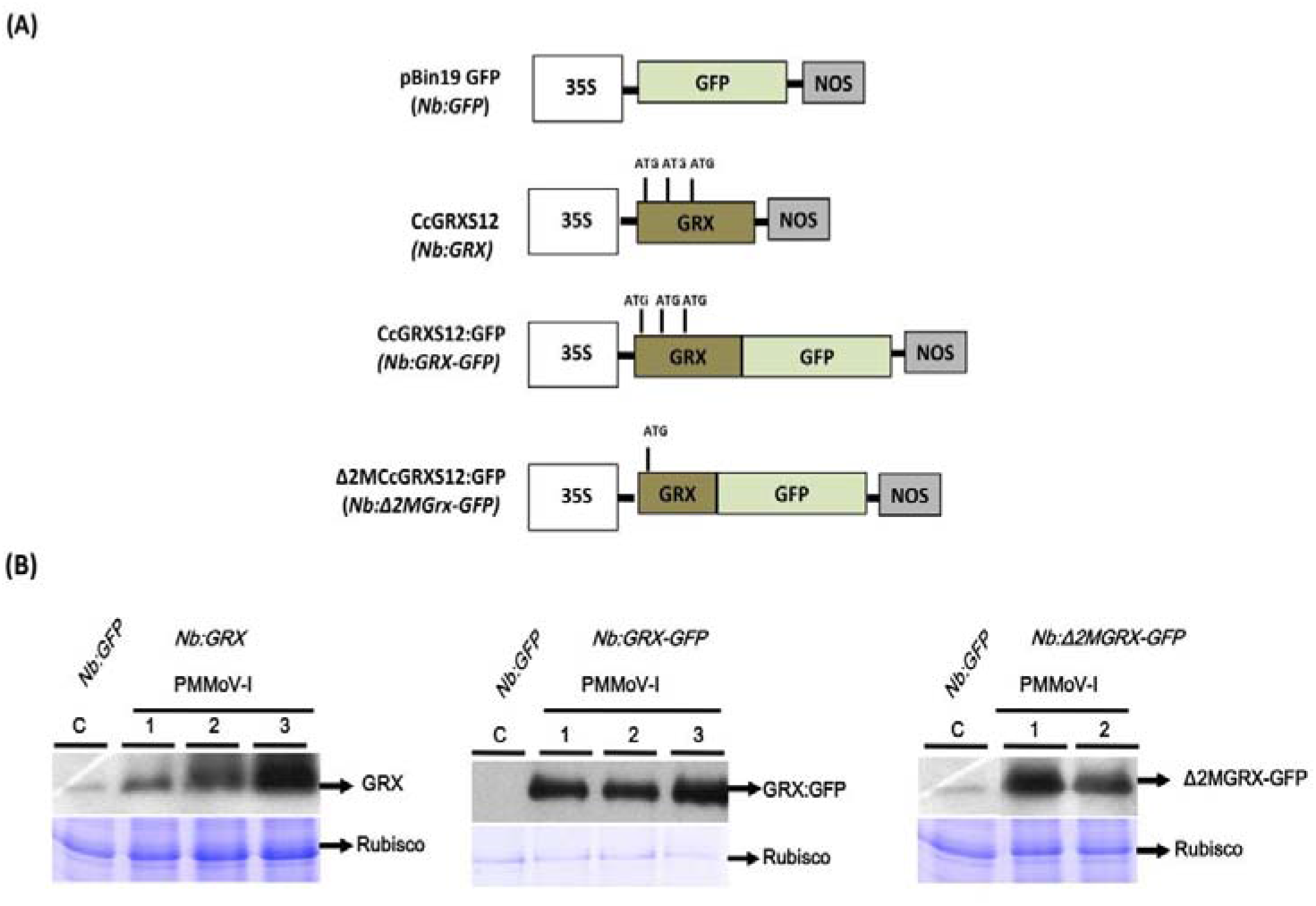
(A) Various gene constructs harboring GRXs used for the transformation of *N.benthamiana* plants and the resultant transgenic lines are mentioned in the parenthesis. (B) Western blot analysis of CcGRXS12 expression in the upper leaves of *N. benthamiana* infected transgenic lines. The protein extracts from *Nb:GRX-GFP* are diluted 10 times when compared with other lines. Lower panels show coommassie staining of total proteins as loading control.

### 3.4. Sub-cellular localization of *CcGRXS12*

To investigate the *in vivo* sub-cellular localization of *CcGRXS12*, expression of *GFP*-fused *CcGRXS12* gene was analyzed in the protoplasts of different transgenic lines *viz*., *Nb:GFP*, *Nb:GRX-GFP* and *Nb*:Δ*2MGRX-GFP* utilizing confocal microscopy. The fluorescence of the native GRX-fused GFP superimposed with chloroplast while the free GFP expressing line (*Nb:GFP*) and the line over-expressing the truncated form of the GRX fused GFP (*Nb*:Δ*2MGRX-GFP*) showed expression signal throughout the cytoplasm and also in the nuclei (Fig.3A). These results confirmed that CcGRXS12 is targeted to the chloroplast and the N-terminus amino acids (63 in number) are essential for the protein to get localized in chloroplast. Western blot analysis with GRX- and GFP-specific antibodies shows that the GRX expression in the transgenic lines are GFP fused one (Fig. 3B).

**Fig.3.**
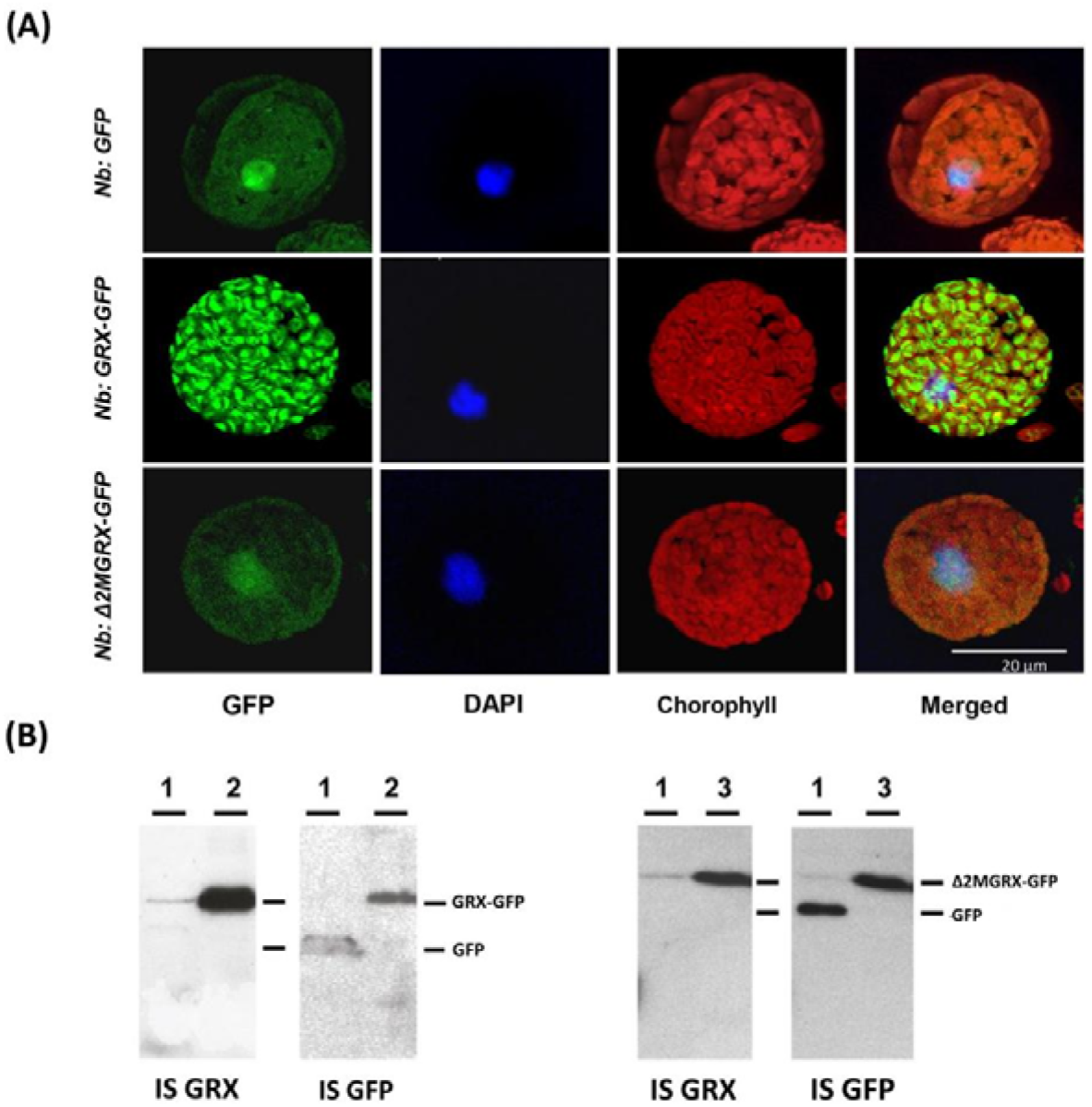
(A) Confocal microscopic study of *CcGRXS12* in *N.benthamiana* protoplast obtained from different transgenics. The protoplasts obtained from different transgenic lines expressing *GFP* are used for analysis: *Nb:GFP ; Nb:GRX-GFP; Nb*:Δ*2MGRX-GFP*. (B) Western blot analysis of CcGRXS12 fused GFP and free GFP in the transgenic lines (1. *Nb:GFP;* 2. *Nb: GRX-GFP*; 3. *Nb*:Δ*2MGRX-GFP*). Visualization was performed for GFP flouresence, chlorophyll autoflouresence, and nuclear staining with DAPI.

### 3.5. Over expression of *CcGRXS12* inhibits PMMoV-I accumulation

A time-course study (7, 14 and 28 dpi) on viral coat protein (CP) titres in the control and *GRX* over-expressing transgenic lines following viral infection were analyzed utilizing coomassie stained gels. At 7 dpi, no difference in viral CP accumulation was found between the inoculated and systemic leaves and also between the different transgenic lines while at 14 and 28 dpi *CcGRXS12* over-expressing lines showed reduced accumulation of viral CP compared with control plants that were transformed with vector devoid of GRX. At 28 dpi, transgenic lines over-expressing *CcGRXS12* showed a significant reduction in viral CP accumulation compared to the GFP expressing lines (Fig.4A). Northern blot analysis for viral gRNA accumulation showed that at early stage of infection (7 dpi), no difference exists between the systemic and inoculated leaves and also between different transgenic lines (Fig.4B). However, at the late stage of infection (28 dpi), the transcript level of viral gRNA in the *Nb:GFP* infected plants were relatively high than other *CcGRXS12* over-expressing lines. The relative ratio of viral RNA accumulation in the *Nb:GRX*, *Nb:GRX-GFP* and *Nb:GRX-GFP* were 0.65, 0.05 and 0.11 respectively when the value was set at 1.0 for the *Nb:GFP* infected lines (Fig.4B). It demonstrates that over-expression of *CcGRXS12* is not inhibiting virus accumulation at early stage but it severely attenuates the accumulation of viral nucleic acids during the later stages of infection.

**Fig.4.**
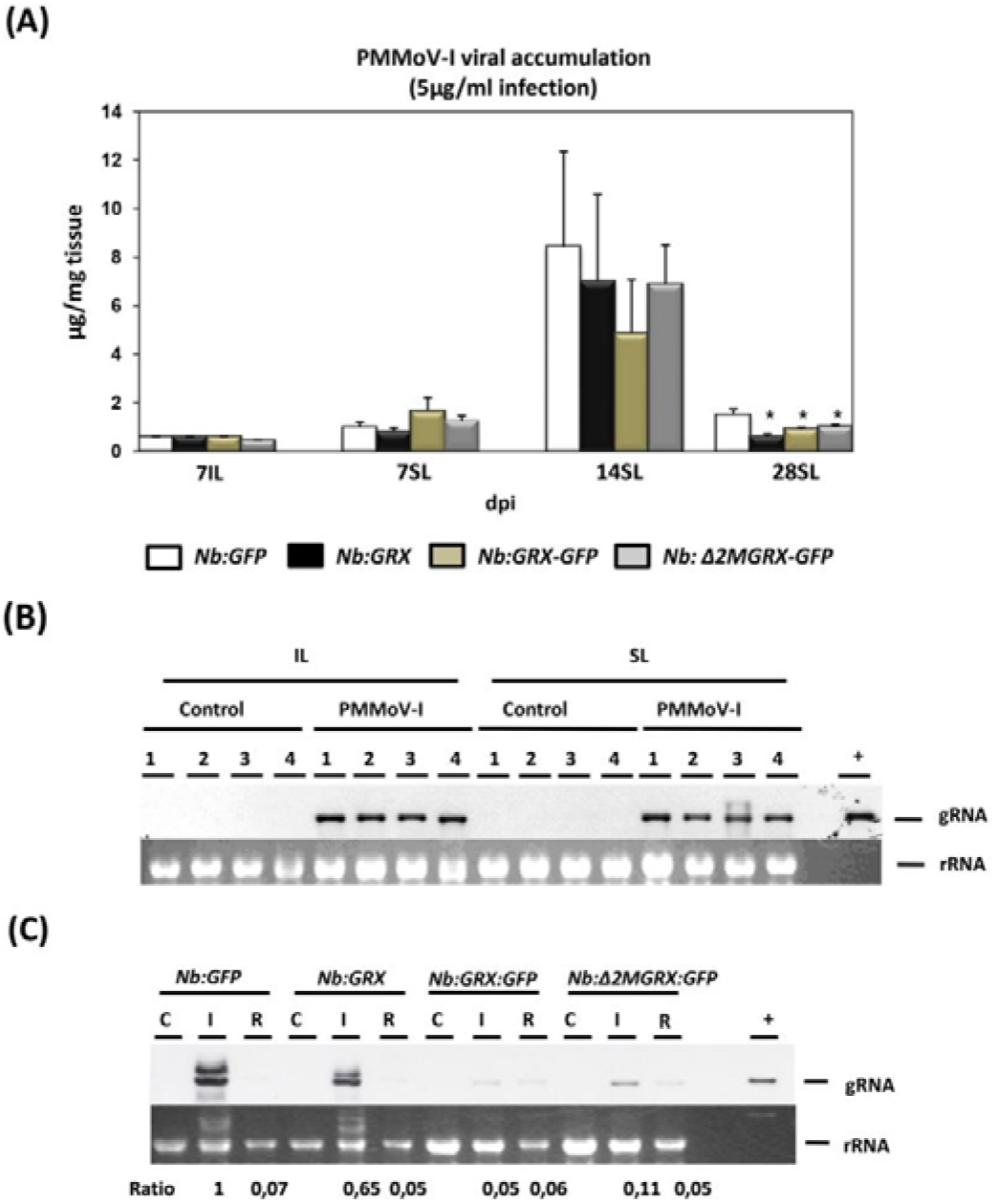
(A) PMMoV-I coat protein accumulation at different time periods (7,14, & 28 dpi) in different transgenic lines of *N. benthamiana*. IL-Inoculated leaves; SL- Systemic leaves. Results are the median of three experiments and are expressed as μg of virus per mg of fresh tissue. Standard deviation are shown and significant differences with respect to *Nb:GFP* are indicated by asterisks (p<0.05). Northern blot analysis of viral RNA accumulation at 7 dpi (B) and 28 dpi (C). The different transgenic lines are represented with numbers. In Fig.B, 1-*Nb:GFP*; 2-*Nb:GRX*; 3-*Nb:GRX-GFP* and 4.*Nb*:Δ*2MGRX-GFP*. The samples analyzed at 28 dpi are marked with alphabets. C-mock control; I- infected; R-recovered leaves. Around 50 ng of RNA extracted from PMMoV-I virus was used as positive control (+). Ratio shown at 28 dpi are the ratio of gRNA accumulation with respect to *Nb:GFP* lines. The lower panels of (B) and (C) are the ribosomal RNA (rRNA) stained with ethidium bromide that served as loading control.

### 3.6. *CcGRXS12* does not inhibit viral RNA replication

In order to study the inhibitory role of *CcGRXS12* towards viral replication, protoplasts obtained from different transgenic lines were infected with viral RNA and the accumulation of the viral CP dynamics was investigated. Viral CP accumulation was ascertained through western blot and it was found to be similar in all lines irrespective of *GFP*- or *CcGRXS12*-expressing lines (Fig.5A). It shows that *CcGRXS12* is not inhibiting the viral RNA replication.

**Fig.5.**
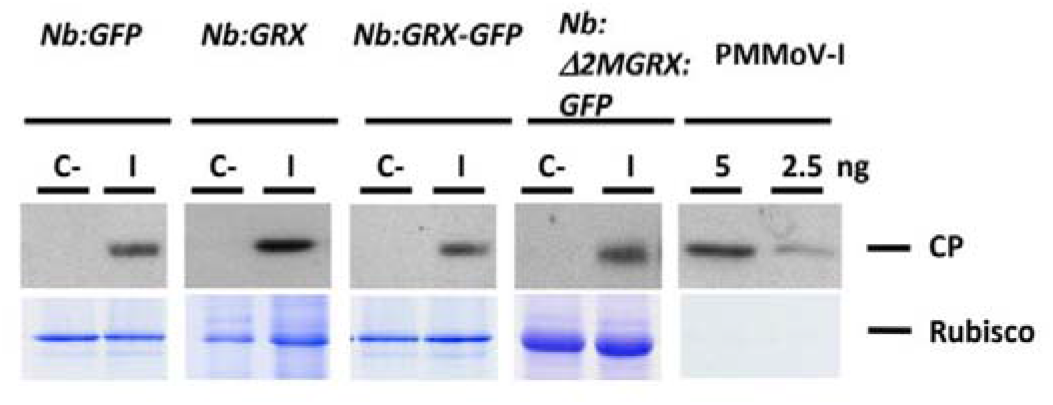
Detection of PMMoV-I coat protein (CP) by western blot using the protoplasts of different transgenic lines. Purified PMMoV-I (5 and 2.5 ng) was used as positive control. C- control uninfected protoplast; I- infected protoplast. Lower panels show Coommassie staining of protein as loading control.

### 3.7. Expression dynamics of transcripts of defence genes

To understand the virus resistant mechanism provoked by *CcGRXS12* in plants, the expression of selective transcripts involved in SA pathway (PR-1, PR-5 and PR-2a), JA/ET pathway (PR-2d) (Van Loon et al., 2006), auxin induced GST marker (pCNT103) (van der Zaal et al., 1987; Droog et al., 1995) were analysed during early (7 dpi) and late stages (28 dpi) of viral infection. At 7 dpi, samples were taken from the inoculated and systemic leaves of the infected plants while at 28 dpi, plants recovered from PMMoV-I infection, thus analysis were done in both the symptomatic and asymptomatic leaves. PMMoV-I -inoculated leaves from *N. tabacum* cv Xanthi (a well-documented PR protein expressing host) (Stintzi et al., 1993; Van Loon et al., 2006), was considered as a positive control for the analysis.

At 7 dpi, high level of SA- pathway related PR transcript accumulation was found in the PMMoV-I- inoculated leaves with no detection in the mock-inoculated leaves, and the level of expression is same irrespective of the transgenic plants (Fig.6A). Expression of basic PR-2d was found in both the mock- and PMMoV-I- inoculated leaves as the mechanical injury induce JA/ET marker genes. Systemic leaves of the PMMoV-I infected plants showed low level of expression while no expression in the mock-inoculated control plants.

**Fig.6.**
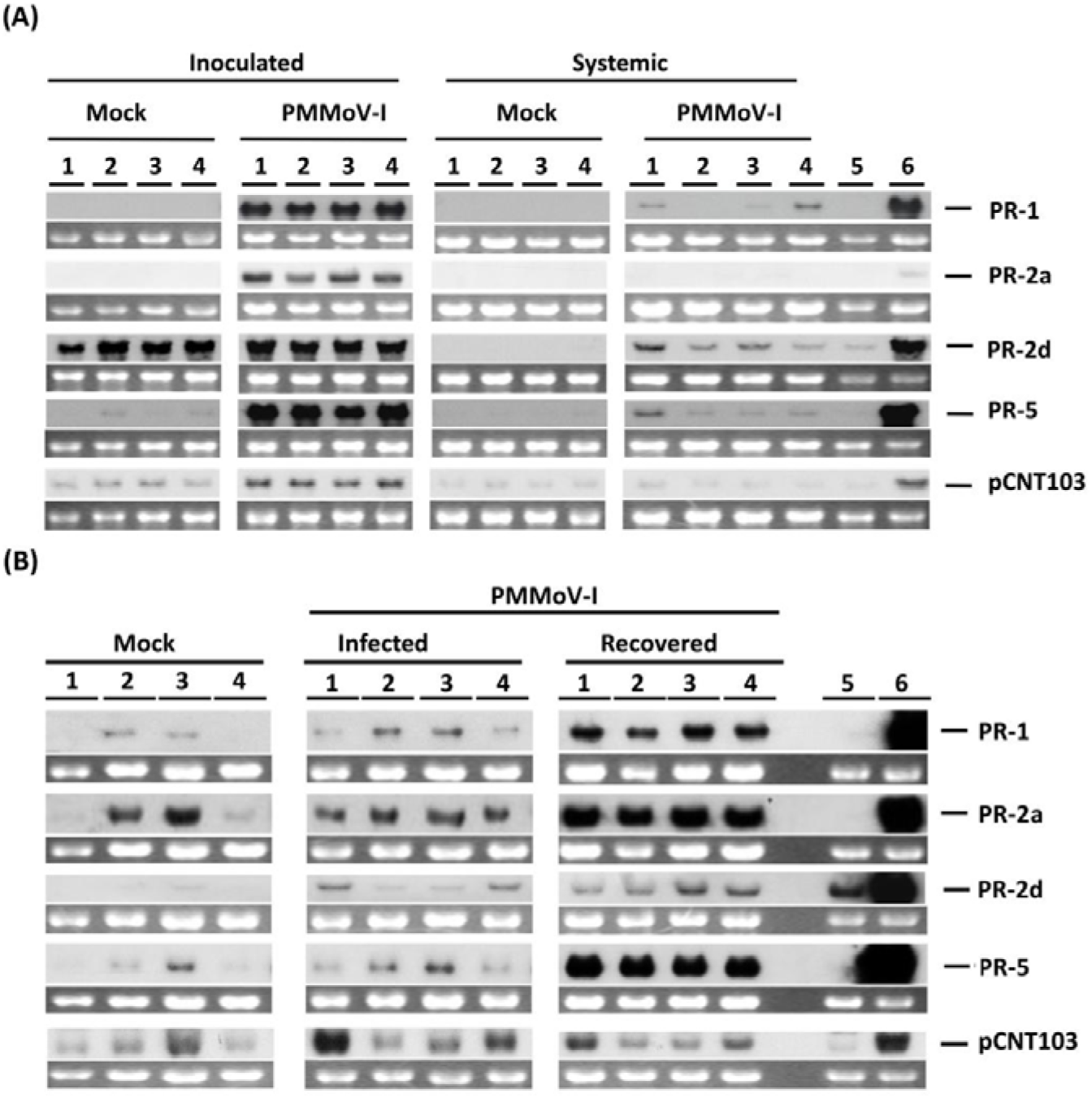
Northern blot analysis of PR transcripts involved in different hormonal pathways (SA, JA**/**ET & auxin) from mock and PMMoV-I infected plants. The expression analysis was performed at 7 dpi (A); and 28 dpi (B). The different transgenic lines are marked with numbers represent: 1-*Nb:GFP*; 2-*Nb:GRX*; 3-*Nb:GRX-GFP* and 4-*Nb*:Δ*2MGRX-GFP*; 5 and 6-mock and PMMoV-I inoculated leaves from *N. tabacum Xanthi* nc plants respectively. The lower panel of each probe shows the ribosomic RNA (rRNA) stained with ethidium bromide. The control (mock) and infected plants are marked above the figure.

At 28 dpi, in the mock-inoculated plants, the accumulation of SA- pathway PR transcripts were found in the *CcGRXS12* over expressing lines that were targeted to the chloroplast. In the symptomatic leaves of the PMMoV-I infected plants, high level of SA- pathway PR transcripts were accumulated in the *Nb:GRX*, and *Nb:GRX-GFP* transgenic lines whereas in the *Nb:GFP* and *Nb*:Δ*2MGRX-GFP* lines a little expression was found. The accumulation of JA/ET pathway (PR-2d) and GST (pCNT103) transcripts were found to be suppressed in the *Nb:GRX*, and *Nb:GRX-GFP* transgenic lines when compared with the *Nb:GFP* and *Nb*:Δ*2MGRX-GFP* lines. In the asymptomatic (recovered) leaves of the PMMoV-I infected plants, the accumulation of SA pathway PR transcripts were found to be very high (2-3 folds), whereas no difference for JA/ET pathway marker transcript was observed when compared to the symptomatic leaves. Between the transgenic lines, no differences in accumulation of transcripts involved in the SA-pathway and JA/ET exist whereas the GST (pCNT103) marker transcript was reduced in the *CcGRXS12* expressing lines when compared to the free *GFP* expressing control line.

### 3.8. Effect of CcGRXS12 over expression on redox carrier molecules

At the late stage of PMMoV-I infection (28 dpi), viral suppression and differences in phytohormonal PR transcript accumulation were observed in the *CcGRXS12* over expressing plants. Redox carrier molecules are known to play significant roles during the pathogen attack and during the induction of defense genes (Pétriacq et al., 2013). Hence the accumulation of redox carrier molecules (oxidized and reduced forms of PNs) were analysed in all the transgenic plants at late stage of infection (28 dpi). The samples analyzed were leaves of mock-control, symptomatic (infected) and asymptomatic (recovered) PMMoV-I infected plants.

In the mock-inoculated control plants, the accumulation level of PNs (NAD(P)/(H)) in the free GFP- and *CcGRXS12*-over expressing lines were similar and no significant difference exist among the different lines. Compared to the mock- control plants, the PMMoV-I infected plants (symptomatic leaves) showed high, yet non-significant, accumulation of oxidized form of PNs (NAD^+^ and NADP^+^). Asymptomatic leaves, showed higher accumulation of (NAD^+^ and NADP^+^) and the accumulation level of NADP^+^ is significantly higher than observed in the mock-control and symptomatic plants.

In the PMMoV-I infected plants, accumulation of the reduced form of PNs (NADH & NADPH) in the symptomatic and asymptomatic leaves were significantly increased compared to mock-control plants. NADH accumulation in the symptomatic leaves was 7-9 times higher than the mock-control plants. The accumulation level of NADH in the *Nb:GRX-GFP* line was found to be significantly high when compared to other lines. Although increase in NADH level was observed in the asymptomatic leaves, the accumulation level was lower than the symptomatic leaves of the infected plants. The NAD pool is increased considerably in the infected plants and it is mainly due to NADH accumulation. Thus, in the PMMoV-I infected plants the NAD pool gets shifted considerably towards its reduced form. Increase in NADPH content was found in the PMMoV-I infected plants (symptomatic and asymptomatic leaves) and symptomatic infected plants showed significantly higher accumulation. As with NADH, the accumulation of NADPH in the infected *Nb:GRX-GFP* plants was significantly higher when compared with other lines. Even though the level of NADPH is increased in the PMMoV-I infected plants, the NADP pool is maintained in the oxidized state.

### 3.9. Role of CcGRXS12 in abiotic stress tolerance

Contribution of *CcGRXS12* to abiotic stress tolerance was analysed by root growth assay. When transgenic lines were grown in the media containing 3 mM H_2_O_2_ or 1 μM paraquat, plants over expressing *CcGRXS12* and its derivatives showed significantly increased primary root elongation than the free *GFP* transgenic line (*P*<0.05) (Fig.11B). The effect of paraquat treatment was stronger than the H_2_O_2_. The abiotic stress tolerance observed was independent of protein localization as *Nb*:Δ*2MGRX-GFP* transgenic lines also show better growth in the oxidative media. No significant differences exist between the free *GFP* and *CcGRXS12* over- expressing plants when grown in the control media. Thus, over expression of *CcGRXS12* in plants increased the abiotic stress tolerance caused by either H_2_O_2_ or paraquat.

### 3.10. Functional substitution of *CcGRXS12* in yeast Δ*grx5* mutants

In yeast, *ScGRX5* was characterised to carry out the Fe-S cluster assembly mechanism (Rodrıguez-Manzaneque et al., 2002). The role of plant GRX in Fe-S cluster assembly mechanism was studied through yeast Δ*grx5* complementation studies (Bandyopadhyay et al., 2008). To examine whether *CcGRXS12* substituted the function role of *ScGRX5* in yeast, yeast Δ*grx5* mutants were transformed with *CcGRXS12*. Analysis has shown that *CcGRXS12* could not rescue the Δ*grx5* mutants in the growth defective media (Fig.9A) and also in the media containing external oxidants (Fig.9B). Further, high level accumulation of free iron (Fig.9C) and low relative aconitase to MDH ratio (Fig.9C&D) in the Δ*grx5* mutants were not restored when *CcGRXS12* was transformed suggesting that *CcGRXS12* could not perform the function for Fe-S cluster assembly in yeast Δ*grx5* mutants.

## 4. Discussion

In this work, we detected the transcript accumulation of a chloroplastic class I GRX gene belonging to S12 subgroup (*CcGRXS12*) when *Capsicum chinense* plants were infected with compatible (PMMoV-I) and incompatible (PMMoV-S) plant virus. The increased accumulation of *CcGRXS12* during the viral infection and also during the cold treatment of the plants (data not shown) has shown that this protein could potentially play vital roles during the viral infection and other abiotic stress conditions. Based on the thermodynamic property analyzed for the PtGRXS12 protein, Eariler studies suggested that this protein has the tendency to accumulate during GSH-mediated mild oxidative stress conditions in plants and also during the glutathionylation process in *Arabidopsis* plants (Courturier et al., (2009a; Dixon et al., 2005; Zaffagnini et al., 2012). PMMoV-I infection and cold treatment induces oxidative stress condition in plants and perturb the GSH redox status in the cells (Hakmaoui, et al., 2012; Kumar et al., 2016). The differential expression of *CcGRXS12* gene during the PMMoV-I infection and cold stress condition in capsicum plants may be attributed to its role in GSH mediated oxidative stress.

CSYS active site of GRX (GRXS12) has been reported in yeast, plants and insect but none from prokaryotes and primitive photosynthetic organisms (*Chlamydomonas reinhardtii*) suggesting its definite functions in the higher organisms. *In vitro* oxidoreductase activity tested for the CcGRXS12 shows that it could reduce the GSH-disulphides formed in the HED assay similar to that of poplar PtGRXS12 (Couturier et al., 2009b), however, the catalytic efficiency of the CcGRXS12 was found to be 10 times less efficient than PtGRXS12 (Fig 1B). The deglutathionylation property possessed by the GRXS12 proteins are important for the regeneration of the antioxidant enzymes such as MSRB1 (Vieira Dos Santos et al., 2007) and PrxII protein (Gama et al., 2008) and helps to maintain the redox poise of the plant cells. In the chloroplast of plant cells, class I GRX (C5/S12) and class II GRXs (S14-S17) co-exist and class I GRXs reduce the substrates by utilizing the reducing equivalents from GSH while class II GRX (GRXS14) utilize reducing equivalents from FTR. Thus the presence of GRXS12 protein has multiple functions in the plant system where GSH plays important role.

### 4.1. *CcGRXS12* role in hormonal pathway activation

As GRXs are abundant in the plant genome, the functional role of the isolated gene was studied in relation to plant-pathogen interaction by over-expressing it in *N.benthamiana* domin plants. Members of the TRX family protein (NtTRXh3) was reported to reduce the multiplication and pathogenicity of TMV and CMV in tobacco plants which was accompanied with the activation of SAR defence related PR genes (Sun et al., 2010). In order to comprehend, if *CcGRXS12* has any molecular role in the activation of systemic acquired resistance (SAR) mechanism, SA-, JA/ET-, and auxin induced GST transcripts, were analyzed in different transgenic plants. *CcGRXS12* over-expressing lines did not show difference in PR transcript accumulation compared to *GFP* expressing plants at the early stage of plant growth or infection (i.e., 7 dpi). However, at 28 dpi, high levels of SA-pathway PR transcripts were accumulated in the mock control lines of *CcGRXS12* targeted to chloroplast (*Nb:GRX* and *Nb:GRX-GFP* lines) (Fig.6B). Although higher accumulation of PR proteins were reported during the senescent stage (Obregon et al., 2001) in tobacco plants, the increased accumulation of SA-regulated PR proteins observed in the *CcGRXS12* transgenic plants appears to be related with the expression of *CcGRXS12* targeted to chloroplast. Moreover, mRNA accumulation corresponding to PR was not detected in the *N. benthamiana* plants expressing free *GFP* at that stage of development. Further the analysis of the defense marker transcripts shows that over-expression of *CcGRXS12* in chloroplast (*Nb:GRX* and *Nb:GRX-GFP* lines) suppress the JA/ET and GST transcript in the PMMoV-I infected lines. Earlier reports have shown that SA pathway activation suppresses the JA responsive genes through the induction of GRXs which interacts with the transcription factors involved in the SA-JA antagonism mechanism (Zander et al., 2012; Wasternack and Hause, 2013). SA also inhibits the auxin response pathway by universally suppressing the auxin-related genes and auxin receptor genes (Wang et al., 2007; Kong et al., 2020). The results described herein are in accordance with those observed for *Arabidopsis* GRX480, that when ectopically expressed in *Arabidopsis* enhanced the expression of SA-inducible marker genes while inhibited the expression of JA-regulated genes (Nadamukong et al., 2007). Viral accumulation was inhibited in the truncated GRX (*Nb*:Δ*2MGRX-GFP*) lines, but the SA- pathway activation and suppression of JA/ET pathway in the infected lines were not observed in these lines. This shows that apart from SAR activation, *CcGRXS12* could use other unknown mechanism for viral inhibition. Activation of SA- pathway related transcripts in the mock control plants; and SA-mediated JA/ET antagonism mechanism found in the infected lines of *Nb:GRX* and *Nb:GRX-GFP* show that the presence of *CcGRXS12* in the chloroplast is necessary for mediating this process as it is not observed in the truncated form of the *CcGRXS12* expressing line (*Nb*:Δ*2MGRX-GFP*) and free *GFP* expressing line (*Nb:GFP*). Quite possibly, *CcGRXS12* may activate the SA-pathway genes either by (i) promoting the SA- biosynthesis inside the chloroplast (or) (ii) affect the redox status of proteins (or) transcription factors involved in the transcription of SA- pathway genes thereby mediating the SA-/JA/ET antagonism in virus infected plants. GRXs are reported to synthesis phytohormones inside the plants. In rice, over expression of the *OsGRX6* increases the cytokinin and gibberellic acid levels in the plants (El-Kereamy et al., 2015; Sharma et al., 2013) by activating the phytohormonal pathway synthesizing genes. Its also noteworthy in *Arabidopsis*, the activity of GCL, a key enzyme involved in the SA biosynthesis in chloroplast is affected by the redox-dependent post-translational modification (Hothorn et al., 2006; Hicks et al.,2007). So it’s of future interest whether *GRXS12* contribute to SA biosynthesis within the chloroplast by modifying the SA biosynthetic pathway genes.

### 4.2. Over-expression of *CcGRXS12* inhibits viral accumulation

When the transgenic plants were infected with PMMoV-I, no difference in viral disease symptom was observed at early stage of infection among the *GFP* and *CcGRXS12* over expressing lines. However, recovery of the plants was more obvious in the *CcGRXS12* over- expressing lines at later stages of infection (28 dpi). At early stage of infection (7 dpi), no difference in the level of virus accumulation between the *CcGRXS12* over-expressing lines and the free *GFP* expressing line, while at the late stage of infection, over-expression of *CcGRXS12* inhibited the viral accumulation when compared to the transgenic *GFP* control (Fig. 4A,B&C). Reduced accumulation of virus in the *Nb:GRX-GFP* expressing line shows that the effect was dose dependent as *CcGRXS12* expression was found to be 10 times higher in this transgenic line (Fig.4C). The viral suppression is independent of *CcGRXS12* localization, as *CcGRXS12* targeted to the cytoplasm also inhibited viral accumulation similar to the lines wherein *CcGRXS12* was targeted to chloroplast. Protoplast infection studies showed that *CcGRXS12* and its derivative expressions are not inhibiting the viral replication. This, data is in variance with the previous reports of Auwerx et al., (2009), where it was shown that the replication of HIV was inhibited by the expression of glutaredoxin-1 in the mammalian cell system. Montes-Casado et al., (2010) have shown that the reduction of PMMoV-I virus accumulation was due to a lower number of infected cells in the systemic leaves of the *CcGRXS12* transgenic plants. Hakmaoui et al., (2012) have shown that tobacco plants infected with PMMoV-I virus recovered at late stage (28 dpi) of infection by up regulating the expression of super oxide dismutase and maintaining the adequate level of peroxiredoxins which are the key antioxidants of the cell. Increased ROS accumulation and decline in antioxidants are prerequisite for the establishment and spread of virus (Clarke et al., 2002; Hakmaoui et al., 2012). During virus invasion, plants use antioxidant machinery to bring down the oxidative stress condition under control. Virus invasion induced the expression of Glutathione-S-Transferase (GST) to control the oxidative stress condition (Chen et al., 2013; Xu et al., 2013; Pavankumar et al., 2017; Skopelitou et al., 2015). At 28 dpi, the expression of auxin induced glutathione-S-transferase (GST) (pCNT103) was lowered in the infected and recovered leaves C*cGRXS12* and its derivatives expressing line while in the free *GFP* expressing lines it was at high level. The reduced accumulation of pCNT103 transcripts in the symptomatic and asymptomatic leaves of *CcGRXS12* expressing lines showed that the protein (CcGRXS12) enhances ROS- scavenging activity in the *CcGRXS12* over expressing line and thus limit the virus induced oxidative stress condition. It’s note worthy to mention that *AtGRXS12* could regenerate PrxII and MSR B antioxidants in Arabidopsis (Vieira Dos Santos et al., 2007 & Gama et al., 2008). Individual GRXs have different regulatory roles on ROS homeostasis. Over expression of *ROXY1* GRX accumulate higher ROS content whereas *ROXY18/GRXS13* over expression reduces the ROS accumulation (Wang et al.,2009; La Camera et al., 2011). Our *in vitro* abiotic stress tolerance assays also shows that *CcGRXS12* over expressing lines are found to be resistant against oxidative stress conditions caused by paraquat and H_2_O_2_. Our combined studies have shown that the protein could able to protect the plants from oxidative stress condition at the time of pathogen attack.

### 4.3. *CcGRXS12* maintains the redox status of the plant cells

Redox status of the cell is sensed and signalled inside the cells by the redox carrier molecules (Ascorbate, Glutathione & Pyridine nucleotides) in oxidized and reduced forms. The inter conversion of the redox carrier molecules between the reduced or oxidized forms as ascorbate (ASC,DHA), glutathione (GSH, GSSG) and pyridine nucleotides (NAD(P)+, NAD(P)H) depends on the cellular redox environment of the cell. *N.benthamiana* plants infected with PMMoV-I virus, shift the redox carrier molecules (ASC,DHA; GSH,GSSG) more towards oxidized condition (Hakamoui et al., 2012). In our work, we have found that PMMoV-I infection of *N. benthamiana* transgenic lines increase the PNs level by shifting PN towards reduced condition as the levels of NADH and NADPH were found to be significantly higher in the PMMoV-I infected plants than in the mock-inoculated control plants at 28 dpi (Fig.7). Increase in PN contents increased during plant’s stress condition improves plants tolerance to oxidative stresses created by biotic and abiotic stresses (Dutilleul et al., 2005; Pétriacq et al., 2012; Ogawa et al., 2016; Awasthi et al., 2019; Zhao et al., 2019). Among the PNs analyzed, increase in NADH level showed 4-8 folds in the PMMoV-I infected plants which shows that NADH is a good marker for PMMoV-I infection. Many studies also imply that increases in NADH content is a mandatory process during biotic and abiotic stress-related defence mechanisms (Ishikawa et al., 2009; Pétriacq et al., 2016, 2012; Ogawa et al., 2009).

**Fig.7.**
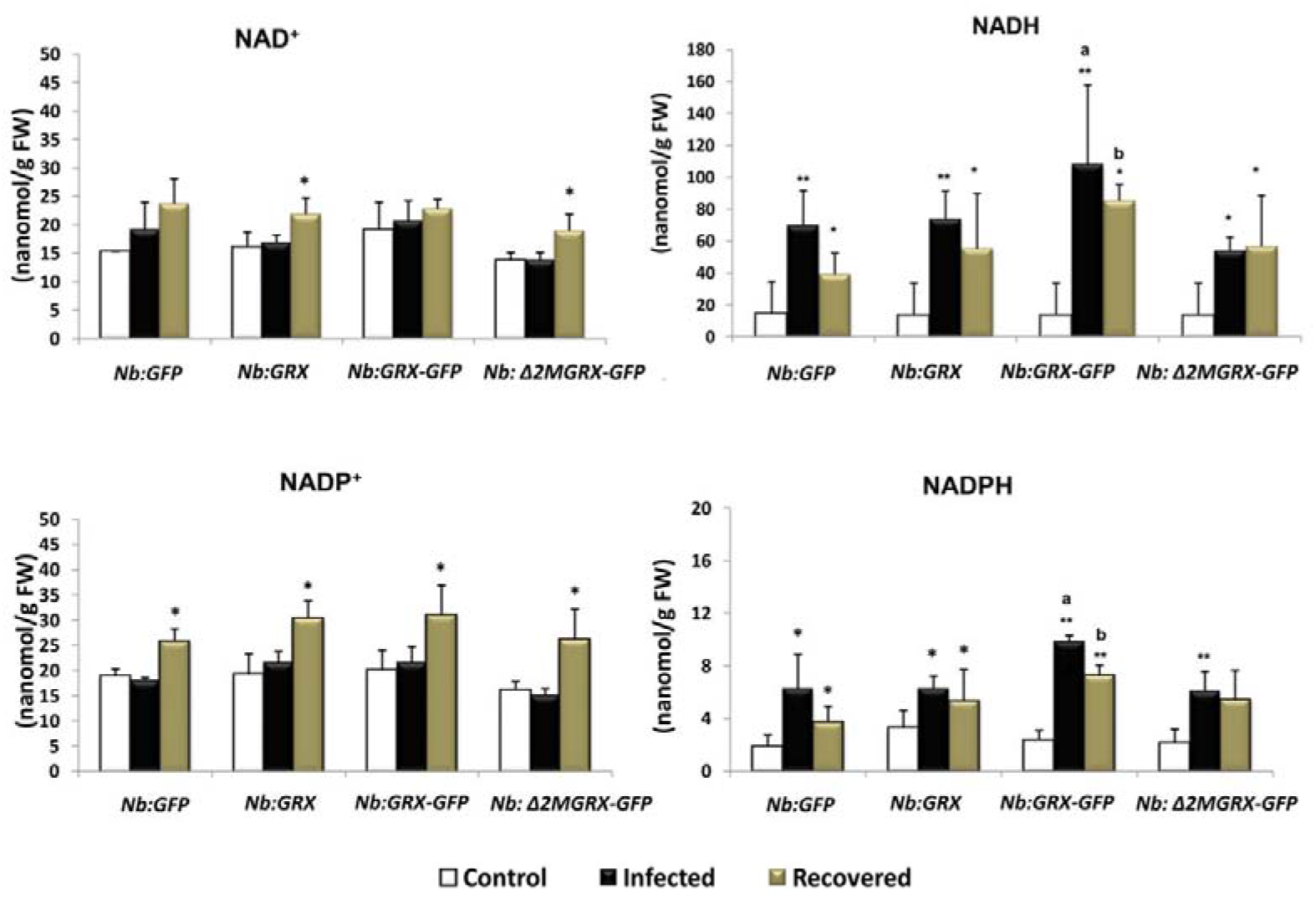
Analysis of pyridine nucleotides (PN) contents in the control, infected and recovered leaves of different transgenic lines. The levels of NAD^+^, NADP^+^, NADH and NADPH were expressed as nanomoles per gram of fresh tissues (nmol**/**g FW). The significant difference between the mock inoculated control and PMMoV-I infected plants is indicated by asterisks (*)significant difference when (p<0.05) and double (**)significant differences when (p<0.001). Significant differences among the different lines in PMMoV-I infected plants and recovered plants is noted by the alphabetical letter ‘a’ and ‘b’ respectively in which the value of p<0.05.

**Fig.8.**
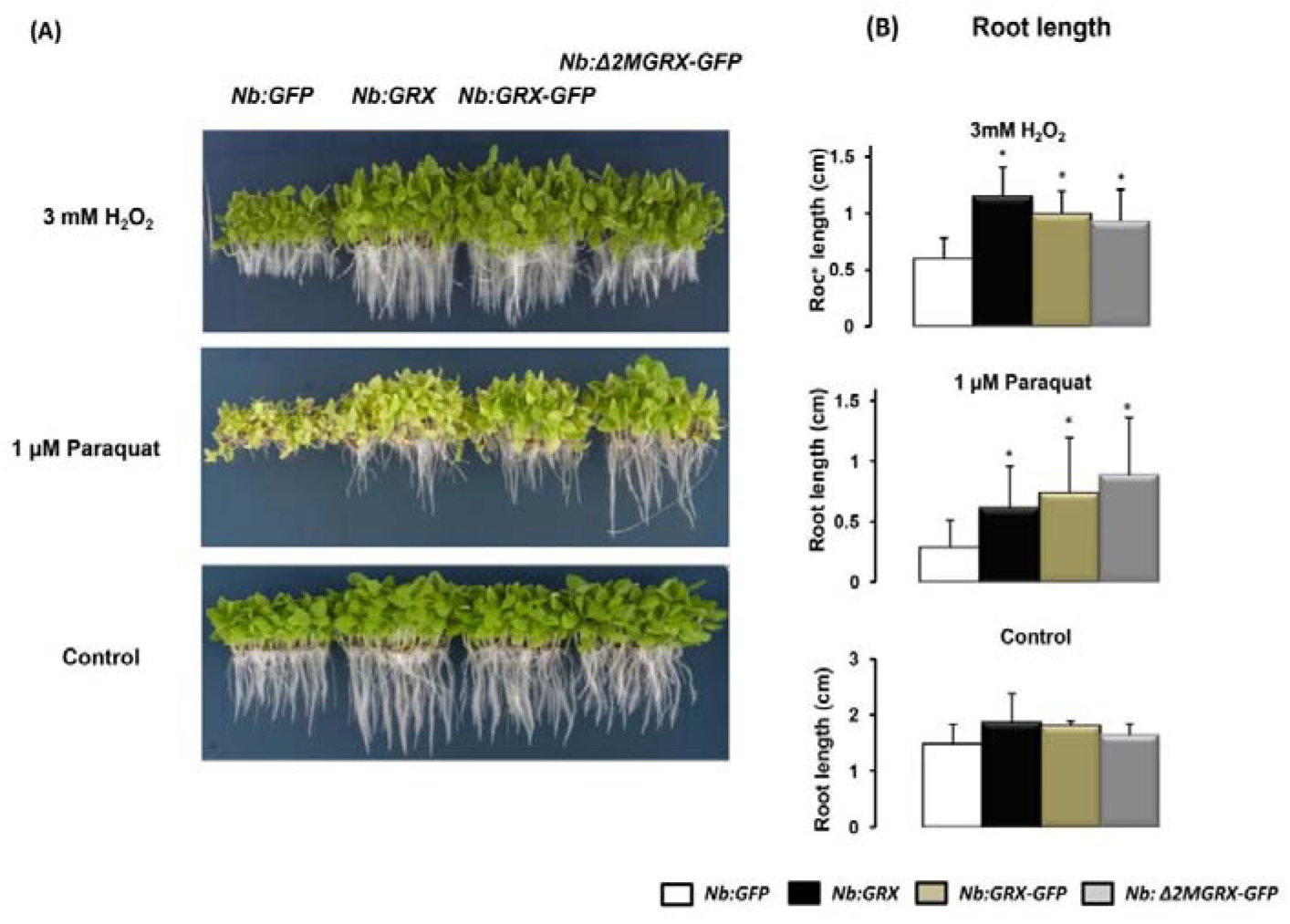
(A) Phenotype of 12 days old *N. benthamiana* transgenic lines grown under normal and oxidative stress conditions (B) Root length measurement of 30 seedlings for triplicate. The length of the primary roots of seedlings was measured in cm. Significant differences between the *Nb:GFP* and *CcGRXS12* over expressing lines are represented by asterisks (*) where p<0.05).

Noctor, (2006) have proposed that reduced form of PNs produced in the cells are utilized by the NAD(P)H consuming enzymes which are involved in the synthesis of ROS and RNS in the cell which may act as a signalling molecule for the plant defence mechanism. Increase in NADH content in the PMMoV-I infected and recovered plant leaves produces an imbalance in the NADH/NAD^+^ that triggers the production of ROS (Millar et al., 2001) or the regulation of cellular antioxidant systems (Dutilleul et al., 2003). Ogawa et al., (2016) have shown that Arabidopsis KO-*nudx6/7* mutants which accumulates high level of NADH are accompanied with increased biotic and abiotic stress tolerance. Increase in NADH in these mutant lines brings biotic/abiotic stress tolerance through the expression of biotic and abiotic stress responsive genes. The genes positively correlated with the increased NADH level belongs to SA pathway (PR1, PR5), JA/ET pathway genes (PDF1.2), oxidoreductase and post-translational modifying (PTM) enzymes. The genes activated by the increased NADH content differed from H_2_O_2_ pathway mediated gene expressions. In our work, we found that within the infected transgenic lines, only *Nb:GRX-GFP* line showed significantly higher NADH and NADPH accumulation at 28 dpi which was accompanied with enhanced virus resistance (Fig.15) which shows that high level *CcGRXS12* expression respond to pathogen infection with increased NADH content.

### 4.4. Redox carrier molecules versus PR proteins

Many studies have correlated the phenomenon of PN accumulation and PR gene expression. Extracellular application of pyridine nucleotides induce plant resistance to pathogen (Zhang and Mou, 2009; Alferez et al., 2018; Wang et al., 2016; Sidiq et al., 2021). (Ge et al., 2007; Reducing the level of pyridine nucleotides through genetic manipulation results in the compromisation of SA pathway activation and pathogen resistance (Li et al., 2021). Nevertheless, our study has proven that *CcGRXS12*-mediated PR transcript accumulation is independent of PN accumulation as it was found that PN accumulation in the different transgenic plants and the mock-inoculated control plants are comparable but the accumulation of SA-pathway transcript was high in *Nb:GRX-GFP* line (Fig.6B). Further at 28 dpi, PN accumulation in the asymptomatic leaves were found to be lower than their infected counterparts, which is contrast with PR accumulation. Thus *CcGRXS12*-mediated PR gene expression is independent of PN levels under pathogenic and non-pathogenic conditions.

### 4.5. *CcGRXS12* in abiotic stress tolerance

It has been demonstrated that many plant GRX proteins protect the plants from oxidative stress conditions created during abiotic stress conditions (Wu et al., 2012; Wu et al., 2017). Treatment of plants with paraquat generates ROS in chloroplasts due to auto oxidation of paraquat radicals generated by electrons from the reaction center of PSI (Taiz and Zeiger, 2010; Krieger-Liszkay et al., 2011), thereby inducing oxidative damage. *CcGRXS12* over expression enhances the plants tolerance against oxidative stress conditions caused by H_2_O_2_ and paraquat irrespective of the protein localization. Different mechanism of GRX- mediated ROS scavenging mechanisms are proposed; GRXs detoxify ROS toxicity through lowering the superoxide ion radicals accumulation (Laporte et al., 2012; Ning et al., 2018) or by regenerating the antioxidant proteins (Rouhier et al., 2005; Wu et al., 2012; Guo et al., 2010; Sharma et al., 2013; Morita et al., 2015). Apart from this, GRXs are reported to scavenge ROS through interaction with transcription factors involved in the stress- related genes expression (Hu et al., 2015; Wu et al., 2012) or by protecting the thiol groups on the enzymes (Morita et al., 2015). The mechanisms through which *CcGRXS12* induce abiotic stress tolerance in *N. benthamiana* have not been analyzed in this work, however it is plausible that the over-expression of *CcGRXS12* could activate the expression of genes involved in the antioxidant mechanisms either in the cytoplasm or the chloroplast. Earlier, *in vitro* studies have reported that *AtGRXS12* could able to regenerate PrxII and MSR B which are the major antioxidants in plant system (Vieira Dos Santos et al., 2007 & Gama et al., 2008).

### 4.6. *CcGRXS12* in Fe-S cluster assembly mechanism

Fe-S cluster containing proteins were found to be abundant in the chloroplast and mitochondria and also throughout the cell (Przybyla-Toscano et al., 2018). Involvement of GRX5 protein in Fe-S cluster assembly was first studied in yeast where the deletion of *GRX5* caused deficient in the synthesis of Fe-S cluster containing proteins and further leads to the accumulation of iron which increases the sensitivity of yeast cells towards oxidative stress conditions (Rodriguez-Manzaneque et al., 1999, 2002). Many plant *GRX*s substitute the function of *GRX5* in yeast (Cheng et al., 2006; Cheng, 2008; Bandyopadhyay et al., 2008). In our experiment, the over expression of *CcGRXS12* in yeast Δ*grx5* mutant could neither restore the Fe-S enzyme activities nor suppress iron accumulation (Fig 9 C & D). Thus, *CcGRXS12* could not participate in Fe-S cluster assembly in yeast but in plants it’ is still uncertain. The structural analysis of populous PtGrxS12 the closest paralog of CcGRXS12 shows that the presence of Trp at −1 position prevents the protein from Fe-S cluster assembly (Couturier et al., 2009b). However, the later studies on AtGRXC5 structure showed that apart from Trp at – 1 position, amino acids at other sites are the deciding factors for the Fe-S cluster assembly mechanism (Couturier et al., 2011). Thus yeast transformation studies showed that the protein may not participate in the biogenesis of Fe-S cluster assembly or in the regulation of iron homeostasis in the chloroplasts.

**Fig.9.**
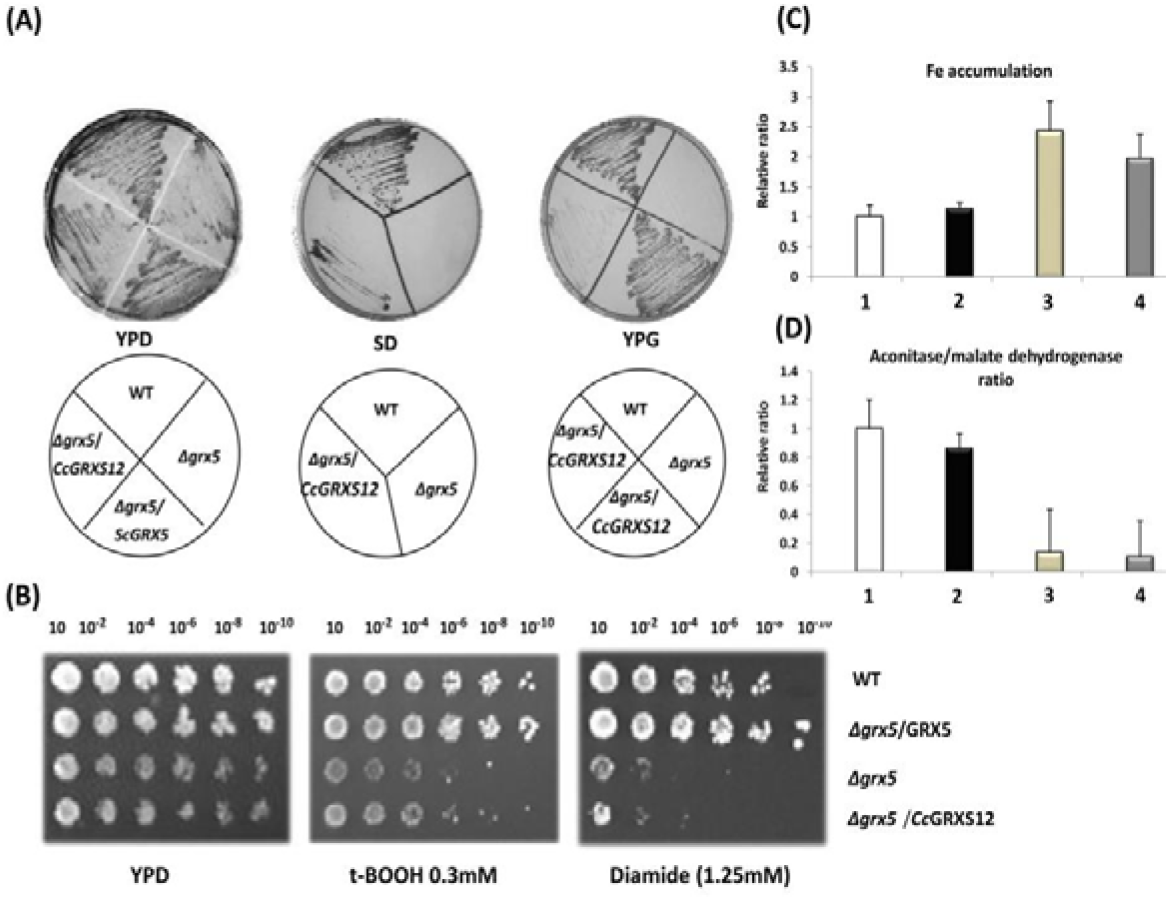
*CcGRXS12* complementation assay in yeast Δ*grx5* mutants. (A) Analysis of the rescue effect of *CcGRXS12* in defective media. The different yeast strains (WT- wild type; Δ*grx5*; Δ*grx5/GRX5*; Δ*grx5/CcGRXS12*) were grown in the glucose (YPD), minimal (SD) and Glycerol (YPG) media for 3 days at 30°C. (B) Sensitivity towards oxidants were analyzed by grown over YPD media containing t-BOOH and diamide for 3 days at 30°C. (C) Relative accumulation of free iron in the different yeast strains. (D) Relative ratio of aconitase to malate dehydrogenase in different yeast strains. In C & D, 1, 2, 3, 4 represent yeast strains: wild type, Δ*grx5*, Δ*grx5/GRX5*; Δ*grx5/CcGRXS12*, respectively.

In conclusion over expression of *CcGRXS12* in *N.benthamiana* plants protect the plants from the oxidative stress conditions created during the biotic and abiotic stresses. CcGRXS12 protein possesses oxidoreductase activity as it could reduce the disulfide bonds formed between GSH and substrate during the HED assay but not able to participate in Fe-S cluster assembly mechanism. The involvement of this protein in multiple stresses warrants further investigation so that it could be exploited for engineering crops with improved stress tolerance.

## Author Contributions

RMS conceived,designed and performed the experiment; wrote the article SVR revised the manuscript and offered critical comments. ST, ZS, AKB contributed in revising the manuscript.

## Acknowledgements

My first thanks to Dr. Maite Serra who was mentor for RMS and also providing me space for conducting doctoral research studies. I thank Dr. Isabel Garcia Luque for her support and knowledge during my research studies. We thank Dr.Enrique for providing the yeast strains and also carry out yeast expression studies. I personally thank Dr. Victoriana Palpuestra for helping me to analyze pyridine nucleotide.

